# Differential reconfiguration of the RNA-interactome during low glucose stress in memory and cytotoxic T cells

**DOI:** 10.64898/2026.06.07.730558

**Authors:** Thomas C.J. Tan, Christos Spanos, David Tollervey

## Abstract

Cellular adaptation to nutrient fluctuation is a fundamental biological process, crucial to cell fate, function, and survival. The choice between aerobic glycolysis and oxidative phosphorylation does not merely satisfy energetic requirements but actively shapes the cellular stress responses. Here we report that during the complex metabolic programming of CD8+ T cells, glucose utilization pathways correlate with the speed and nature of the response to glucose withdrawal. By quantitating systematic RNA-protein interactions in response to glucose withdrawal, we found that effector T cells mount acute transcriptional and post-transcriptional responses to the stress, while memory T cells exhibit slower, more limited responses. The functional dichotomy observed in T cells - between highly glycolytic cytotoxic effector cells and respiratory memory cells - exemplifies how distinct glucose utilization pathways impact immunological fate and function. Understanding the intricate interplay between metabolic modality, glucose pathways, and post-transcriptional control is crucial for deciphering environment adaptations and developing interventions in contexts ranging from immunotherapy to cancer biology.

## INTRODUCTION

All living cells must sense and respond to changes in their environment, with glucose availability being a common primary metabolic cue. The metabolic state of a cell profoundly affects how it handles the stress of nutrient deprivation: whether it predominately utilizes the rapid, but inefficient (∼2 ATP per glucose molecule) process of glycolysis or the slower, but more efficient (30-32 ATP per glucose molecule) process of oxidative phosphorylation.

The phenomenon of “diauxie” or diauxic shift was first observed in *E.coli* ^1^. Available glucose is rapidly metabolized only via glycolysis, even in presence of oxygen, in a process termed aerobic glycolysis. When environmental glucose has largely been depleted, ATP generation is switched to oxidative phosphorylation, using other carbon sources, including the products of fermentation. This metabolic switch was observed in many other organisms, notably including budding yeast and mammalian cells ^2,3^. More generally, aerobic fermentation has been reported in many systems that require rapid energy availability. Mammalian cells that utilize aerobic fermentation include skeletal muscles, activated T cells and cancer cells, where it is termed the Warburg effect.

T cell activation requires a massive metabolic overhaul. A metabolic shift is seen from quiescent naïve T-cells, predominately utilizing oxidative phosphorylation, to an activated “effector” state predominately utilizing glycolysis through aerobic formation. Upon activation, T cells upregulate the transcription factor Myc, which drives the expression of glycolytic enzymes and glucose transporters ^4^. This metabolic reprogramming is further supported by PDK1 and mTOR, which integrate signals to promote HIF-1α activity and aerobic glycolysis ^5^. Effector T cells become highly dependent on this glycolytic flux to fuel their rapid proliferation, and specialized functions such as cytokine production ^6,7^. Conversely, naïve and memory T cells primarily utilize oxidative phosphorylation (OxPhos) and fatty acid oxidation (FAO) ^8^.

Effector cells exhibit less active, “fissioned” mitochondria, reflecting their prioritization of glycolysis, whereas Memory cells are characterized by “fused” mitochondrial networks that are highly efficient at generating ATP through OxPhos ^9^. This activity allows a high level of spare respiratory capacity (SRC), a measure of the ability of mitochondria to increase energy production under stress ^9,10^. This high SRC is a hallmark of memory T cells and is essential for their long-term survival and rapid recall ability upon re-infection ^10^. Because OxPhos generates significantly more ATP (30-32) per molecule of glucose than glycolysis (2), cells with high respiratory capacity are less dependent on continual glucose uptake ^6^.

In cancer models, cells with mitochondrial respiratory defects are hypersensitive to glycolytic inhibitors, whereas cells with fully functional mitochondria can better withstand glucose deprivation by utilizing alternative energy sources ^11,12^. The ability to switch to OxPhos or utilize FAO provides a metabolic safety net that avoids the dramatic drop in ATP that can arise in glycolytic cells during sugar-depletion stress ^12,13^.

The cellular response to glucose depletion involves a sophisticated shift in post-transcriptional regulation. A key example is the role of glycolytic enzymes as RNA-binding proteins (RBPs). In effector T cells, the enzyme GAPDH serves as a bridge between metabolic flux and protein synthesis ^7^. When glucose is abundant and glycolytic flux is high, GAPDH is occupied by its role in the glycolytic pathway. However, when glucose levels drop or glycolysis is inhibited, GAPDH is freed from its metabolic duties and binds to the 3’ untranslated region (UTR) of specific mRNAs, such as *Ifng*. This binding inhibits the translation of the mRNA, effectively shutting down the production of effector cytokines in response to metabolic stress. This mechanism ensures that the cell does not expend resources on specialized functions during conditions of energy depletion. Beyond specific RNA-protein interactions, glucose withdrawal triggers a global inhibition of mRNA translation to conserve energy. This shutdown is mediated by multiple integrated pathways including mTORC1 inactivation ^14^, eIF2α phosphorylation ^15^, reduced intracellular macromolecular diffusion ^16^, and formation of stress granules and other mRNP condensates ^17^. In budding yeast, dissociation of key translation initiation factors from mRNA is evident 30 sec after glucose withdrawal ^18^. In mammalian cells, it remains unclear how the post-transcriptional response to glucose depletion differs between cells utilizing either glycolysis or OxPhos, given their distinct sensitivities to environmental glucose levels.

Here we use primary mouse CD8 T cells, which can be induced to adopt either aerobic glycolysis or OxPhos as their primary metabolic state by growth *ex vivo* in different cytokine environments. Over a time-course following glucose withdrawal from the growth media, RNA-protein interactions were fixed by UV-crosslinking. The total RNA-bound proteome was purified and identified by mass spectrometry.

## RESULTS

### CD8 T cells show distinct responses to glucose withdrawal depending on differentiation state

To identify hallmarks of RNA-protein interactions during low glucose stress we generated T cells preferentially utilizing either OxPhos or aerobic glycolysis for ATP production. Ex vivo primary CD8 T cells were activated for two days, expanded in IL-2 for another two days, and differentiation was induced for three days in SILAC media containing either IL-15 (memory cells, OxPhos, abbreviated as IL15 cells thereafter) or IL-2+IL-12 (cytotoxic effector cells, glycolytic, abbreviated as IL12 cells thereafter). On the day of harvest, cells were transferred to Glu+ (normal) or Glu-(glucose withdrawn) conditional media for set durations, followed by UV-crosslinking and snap-freezing on dry ice. Total cellular ribonucleoproteins were extracted, and RNA-interactomes were quantified using TRAPP-DIA-SILAC, as previously reported ^19^. SILAC data was processed with DIA-NN (v1.8.1) using a gas-phase fractionated DIA spectral library, and analyzed using the workflow shown in Fig S1.

Normalisation of the RNA-protein interactome dataset was performed at the peptide level, using the median of each sample. Proteins represented by more than 2 unique peptides were selected for further analyses, and protein abundance was defined as the median value of all represented peptides (Fig S2). For each protein, recovery was computed as a Glu-: Glu+ ratio. To allow comparison of ratios between time points, the maximal allowed missing value of an individual protein was set to 2 out of 4 replicates in any condition at any time point. These were termed “consistently identified proteins” (Fig S3). A list of RNA-binding proteins in CD8 T cells was constructed using a label-free DIA TRAPP dataset comparing IL15/IL12 cells with/without UV-crosslinking (Fig S4). Intensities were normalized using VSN at the protein level, and proteins with intensities significantly elevated in UV-crosslinked cells (two-tailed T Test with Benjamini Hochberg correction, FDR < 0.05, FC > 1.5) were considered RNA-binding. The consistently identified proteins were overlayed with the RNA-binding proteins, and the common proteins between the sets were used for downstream analysis. Results are summarized in Fig 1A. 6892 proteins were represented by more than 2 unique peptides in IL15 cells, and 7553 proteins in IL12 cells. These are reduced to 2542 consistently identified proteins in IL15 cells, and 4192 in IL12 cells. Of those, 1277 proteins were considered RNA-binding in IL15 cells, and 1556 in IL12 cells.

**Fig 1.**
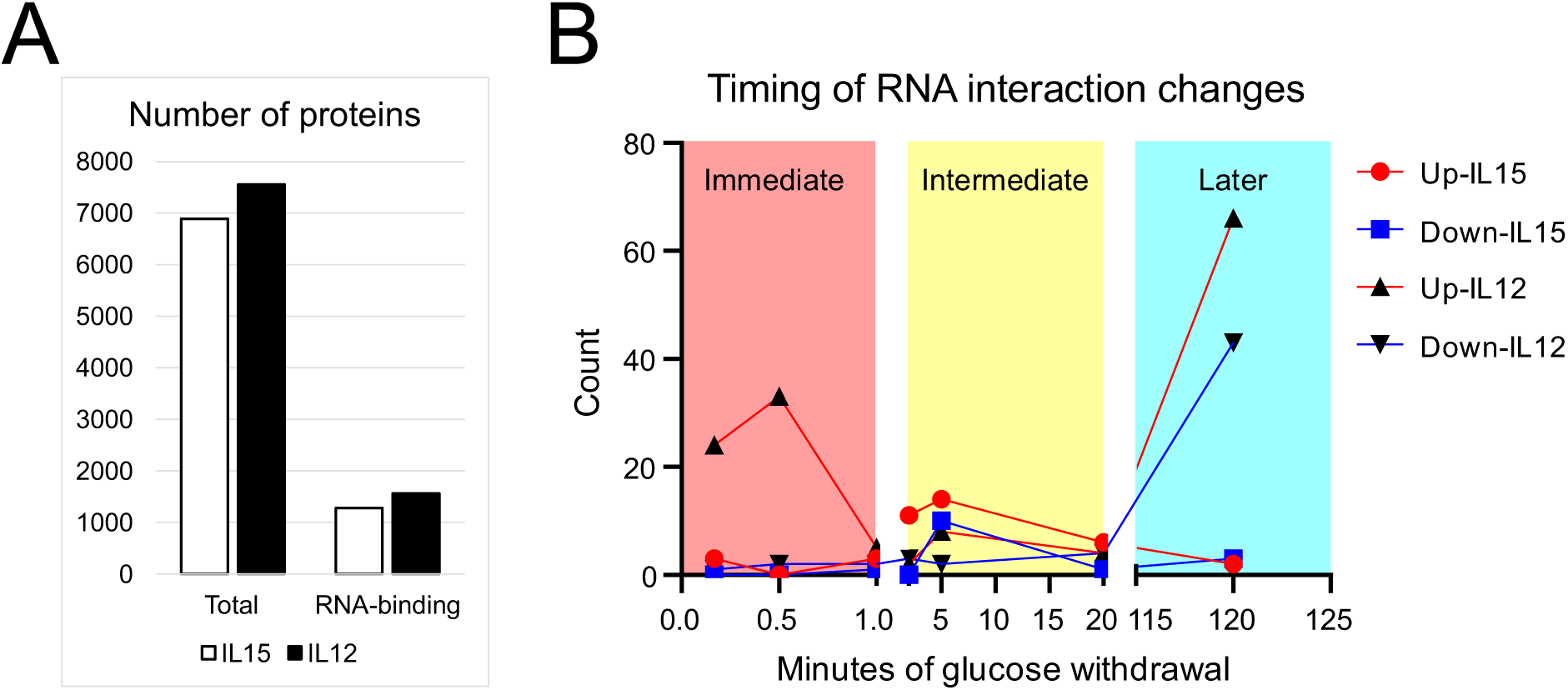
Detected proteins: Timing of RNA interaction changes. (A) Number of proteins passing detection thresholds: total proteome – 2 unique peptides detected; RNA-interactome – 2 unique peptides, detected in at least 2 out of 4 replicate samples at all time points, enriched by UV-crosslinking. (B) For proteins with significantly altered RNA interaction, number of those reaching their highest fold-change were plotted in each time point. Significance was determined by two-way ANOVA (within cell states) or two-tailed T Test (between cell states) at each time point, and verified manually.

The retained protein sets were tested for significant responses to glucose depletion, using two criteria. To identify significant changes in RNA association in either cell state following glucose withdrawal, protein recovery across timepoints was assessed using two-way ANOVA (P<0.05, FC>95 pct or <5 pct). To identify differential RNA association between the cell states, two-tailed T Tests were used to compare IL15 and IL12 cells at each time point (P<0.05, FC >90 pct or <10 pct). For proteins identified as significantly altered, Glu-:Glu+ recovery ratios across the time course were manually plotted and checked for fold-change and consistency. Numbers of significant proteins at their highest fold-change were plotted across the time course in Fig 1B. It became clear that the response to glucose depletion was greater in the glycolytic cells. In OxPhos IL15 cells, 7 proteins showed peak RNA association change in immediate response (the first minute), 42 in intermediate response (between 2-20 min), and 5 in later response (2 h). In glycolytic IL12 cells, 67 proteins showed peak changes in recovery by one minute, 23 between 2-20 min, and 109 at 2 h.

To identify enriched cellular functions, proteins with significantly altered RNA interactions were further analyzed by gProfiler ^20^. The results are listed in Table 1. The OxPhos IL15 cells had a limited response, mostly during 2-20 min post withdrawal (Fig. 1B). Immediate responses were mainly on proteins implicated in specific gene transcription and RNA transport, with intermediate and later responses on specific transcriptional/translational controls and centrosome dynamics. The glycolytic IL12 cells showed more marked changes, with two major peaks; in the first minute, and then over 2 h (Fig. 1B). Immediate responses mainly affected proteins implicated in mRNA transcription and stability, intermediate responses mainly on mRNA stability and nucleoplasm activity, and later responses mainly on metabolic state, transcription, RNA splicing, RNA / protein degradation, and ribosome functions. Supplementary Note 1 describes a number of selected individual proteins with altered RNA association. Selected, enriched GO terms were analyzed in more depth, as reported in the next sections.

**Table 1.**
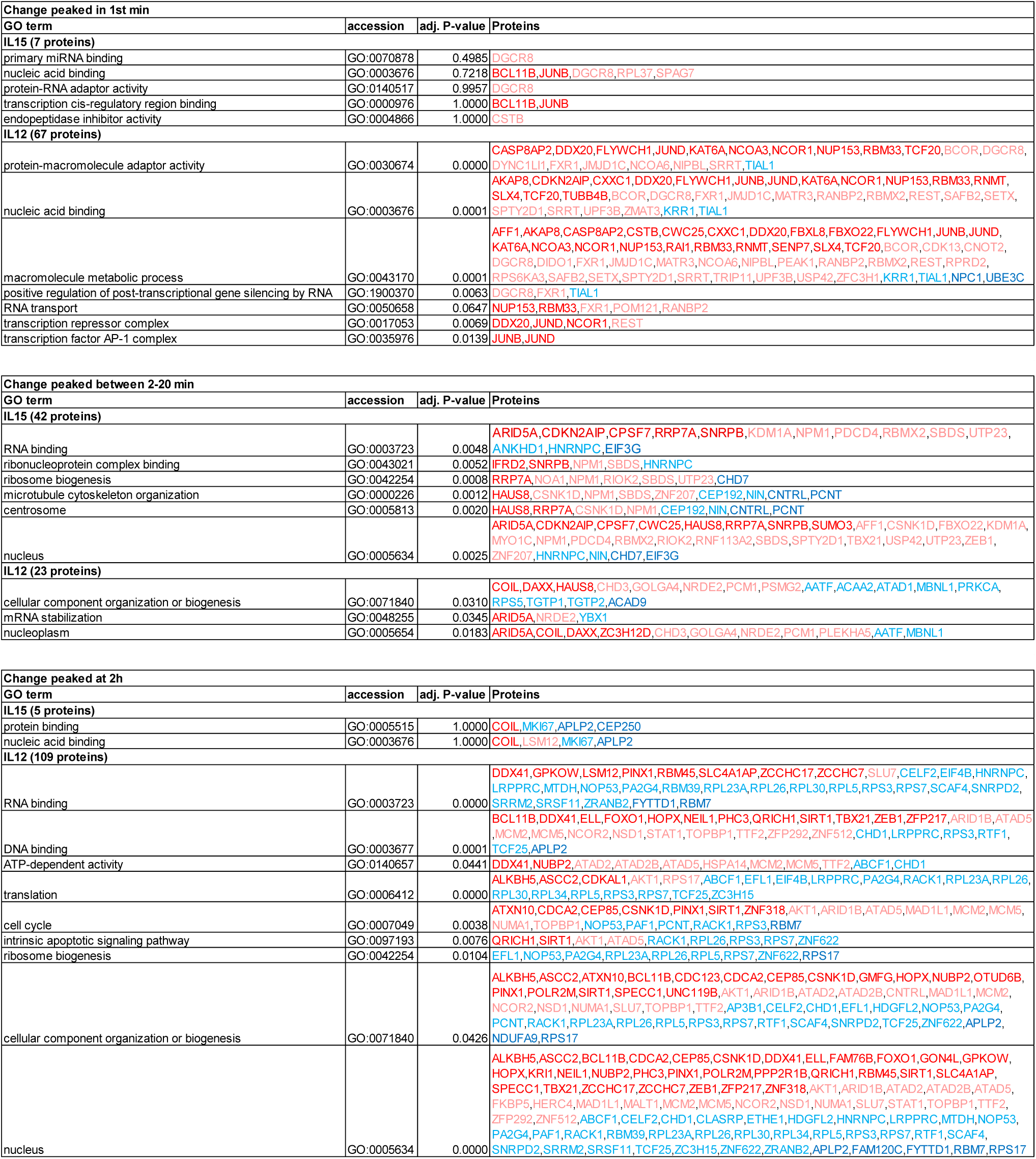
GO term enrichment in altered RNA-interactome. Representative GO terms enriched in proteins with altered RNA-interaction at immediate (first 1 min), intermediate (2-20 min), and late (2 hr) time points following glucose withdrawal in IL15 and IL12 cells. Fold-change in RNA-association are colour-marked: FC≥2 (dark red), 2>FC≥1.5 (pale red), −2<FC≤-1.5 (pale blue), and FC≤-2 (dark blue).

### Translation factors show altered RNA association in IL12 cells

Global mRNA translation shutdown, or altered selection of translated transcripts, constitute the first line of response to cellular stress in other systems ^18,21^ Several ribosomal proteins and translation factors showed reduced RNA association in IL12 cells (Table 1). However, the fold changes were mild and mostly occurred at 2h, indicating a lack of an acute, global translational halt. To analyze these regulators further, Fig. 2A shows a selected set of proteins that are either ribosome components or have regulatory functions in mRNA translation. All are experimentally validated members of GO:0006412 (Biological Process:Translation), plus ribosomal proteins, translation initiation and elongation factors. Fig. 2B highlights ATP-dependent RNA helicases (members of GO:0003724; Molecular Function: ATP-dependent RNA helicase activity).

**Fig 2.**
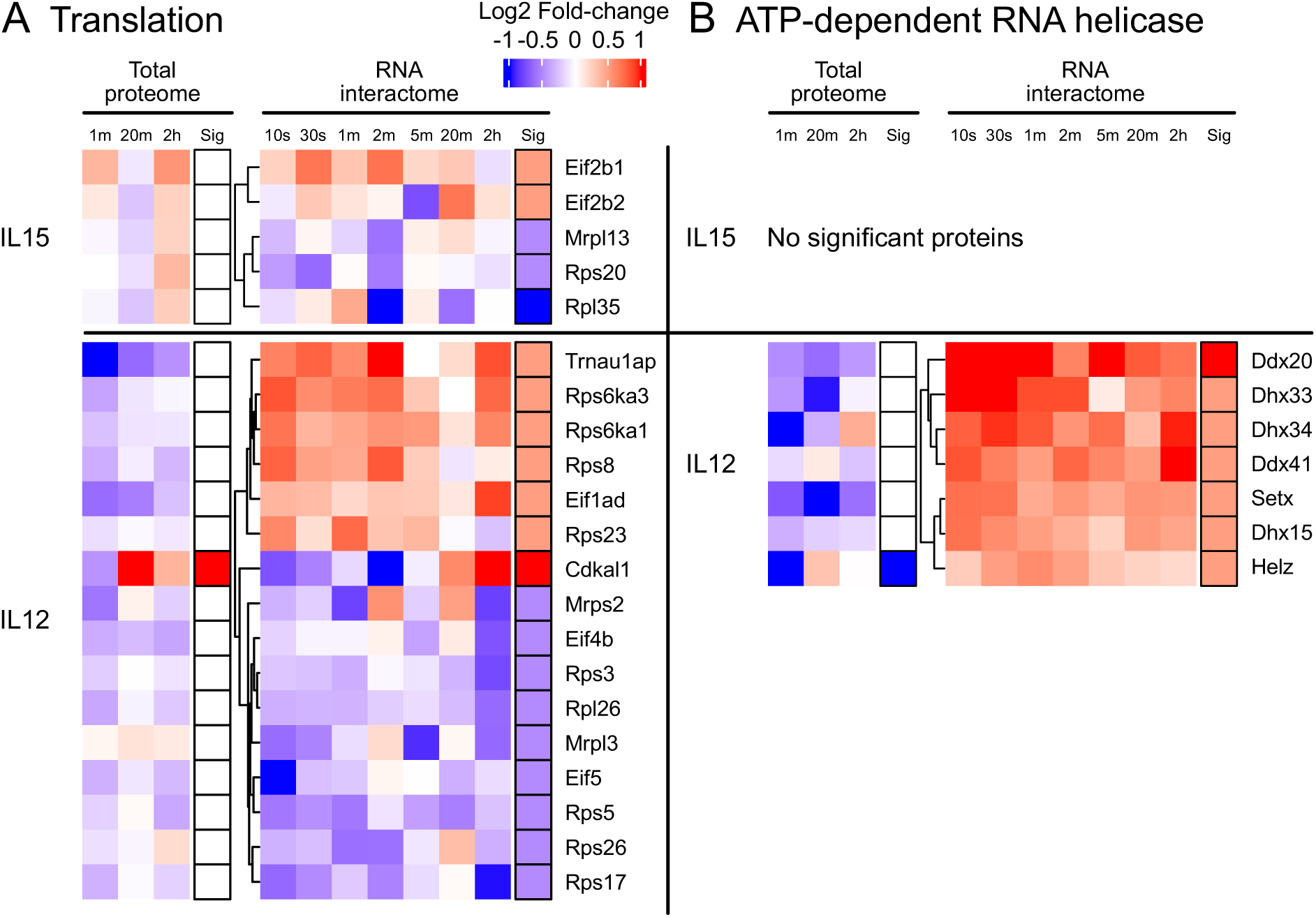
mRNA translation machinery and regulators significantly changed in RNA association following glucose withdrawal. Heatmap showing log2 fold change, from Glu+ to Glu-condition, in IL15 and IL12 cells, across the experimental time course. Clustering was performed based on the RNA-interactome dataset. Statistical significance was determined by the lowest p value in the time course (two-tailed T Test, P<0.05), and the fold change at the corresponding time point (absFC>1.5: pale red/blue; absFC>2: full red/blue). (A) mRNA translation regulators were defined as experimentally validated components of GO:0006412 (Biological Process:Translation), plus all ribosomal proteins, translation initiation and elongation factors. (B) RNA helicases were defined as components of GO:0003724 (Molecular Function: ATP-dependent RNA helicase activity).

Several translation factors displayed consistently altered RNA association upon glucose withdrawal in glycolytic IL12 cells; whereas fewer alterations were seen in OxPhos IL15 cells, and the changes were either transient or inconsistent. Notably, increased recovery of TRNAU1AP, RPS6KA1, RPS6KA3, and RPS8 with RNA was observed at most time points in IL12 cells. Reduced recovery of RPS5, RPS17, and RPS26 could be seen in the first 2 min, while eIF4B, RPS3, RPL26, MRPS2 and MRPL3 were mainly reduced at 2h. TRNAU1AP is required for the production of selenoproteins, and its knockdown affects PI3K-Akt signaling and cell proliferation ^22^. Both RPS6KA1 (RSK1) and RPS6KA3 (RSK2) are p90 ribosomal S6 kinases that act downstream of the ERK/MAPK signaling pathway and regulate the activities of the translation initiation factor eIF4B and the translational repressor PDCD4 ^23^. 40S ribosomal subunit component RPS8 controls translation of specific transcripts ^24^ and is required for cell cycle progression ^25^. RPS5 is located at the mRNA exit channel on 40S ribosomes and regulates translation by direct mRNA interaction ^26^. RPS17 is enriched in nucleolus and may affect pre-ribosome assembly ^27^. RPL26 interacts with p53 mRNA in regulating stress response ^28^. eIF4B plays a key role in loading 43S pre-initiation complexes, together with the helicase activity of eIF4A ^29^. RPS3 directly interacts with mRNA at the entry channel on 40S ribosome, affecting transcript-specific translation and stability ^30,31^. MRPS2 was reported to affect mitochondrial ribosomal protein levels and OxPhos activity ^32^. MRPL3 regulates immune cell infiltration and immune checkpoint gene expression in tumor microenvironment ^33^.

RNA helicases showed altered RNA-association only in IL12 cells, with significantly increased association for DDX20 and DHX33 mainly during the first minute, while DHX34 and DDX41 mainly increased at 2 hr. DDX20 is a multi-functional protein involved in RNA splicing, transcription, and translation control ^34^. DHX33 promotes translation initiation and activates inflammasomes in macrophages ^35,36^. DHX34 is involved in nonsense mediated decay and splicing of pre-mRNAs, leading to negative regulation of T cell immunogenicity in tumor cells ^37,38^. DDX41 functions as a DNA sensor facilitating the type I interferon responses, and also splicing of pre-mRNAs and small nucleolar RNAs affecting ribosome biogenesis ^39^.

Following glucose withdrawal in yeast, the DEAD-box RNA helicases eIF4A and Ded1 are rapidly lost from mRNA 5’ regions, linked to an 80-90% reduction in cellular ATP levels ^13,18^. In contrast, recovery of eIF4A and the Ded1 homolog, DDX3X, were apparently unchanged in both IL15 and IL12 T cells (Fig S5A). In IL12 cells around 30-40% reduction in cellular ATP, from ∼2.4 mM to ∼1.5 mM, was seen after 1 min of glucose depletion, with partial recovery by 2h (Fig S5B). Corresponding reductions were seen for GTP, CTP and UTP, which all require ATP for regeneration (Fig S5B). These represent substantial reductions but would not be predicted to directly induce dissociation of DEAD-box translation regulators, which generally have Michaelis constants (Km) for ATP between 100-500 µM ^40,41^.

### Altered translation regulator abundance following glucose withdrawal

Changes in protein recovery between RNA-interactome datasets could reflect alterations in either its RNA-association or total cellular abundance. Total proteomic datasets were therefore generated from IL15 and IL12 cells, in Glu- and Glu+ conditions at 1 min, 20 min, and 2 hr. The data was acquired as single shot, label free, data-independent acquisition, (DIA) mass-spectrometry (MS). MS results were converted to protein copy number per cell, with a workflow described in Fig S6. Importantly, apart from one instance in IL12 cells (CDKAL1), no proteins classed as showing significant changes in RNA association, also showed clear changes in total abundance in the same direction at the same time point. We conclude that alterations described above genuinely reflect changes in protein-RNA interactions.

In agreement with changes in RNA association, alterations in cellular protein abundance were more prominent and acute in IL12 cells relative to IL15. In IL15 cells, only 27 proteins passed the two significance thresholds at 1 min, 67 at 20 min, and 86 at 2 hr. In comparison, in IL12 cells significant changes were observed for 406 proteins at 1 min, 205 at 20 min, and 180 at 2 hr.

Analyses of proteins with altered cellular abundance initially focused on translation-related factors (Fig 3). We noted changes in enzymes responsible for post-translational modification of translation factors. eEF1A is trimethylated at Lys79 by eEF1AKMT1 (N6AMT2) ^42^, and at Lys318 by eEF1AKMT2 (METTL10) ^43^. eEF2 is trimethylated at Lys525 by eEF2KMT (FAM86A), facilitating productive engagement with ribosome ^44^. In contrast, phosphorylation of eEF2 at Thr56 by eEF2K reduces translation elongation rates ^45,46^. Global translation repression is mediated via eIF2α phosphorylation at ser51 by multiple eIF2AKs.

**Fig 3.**
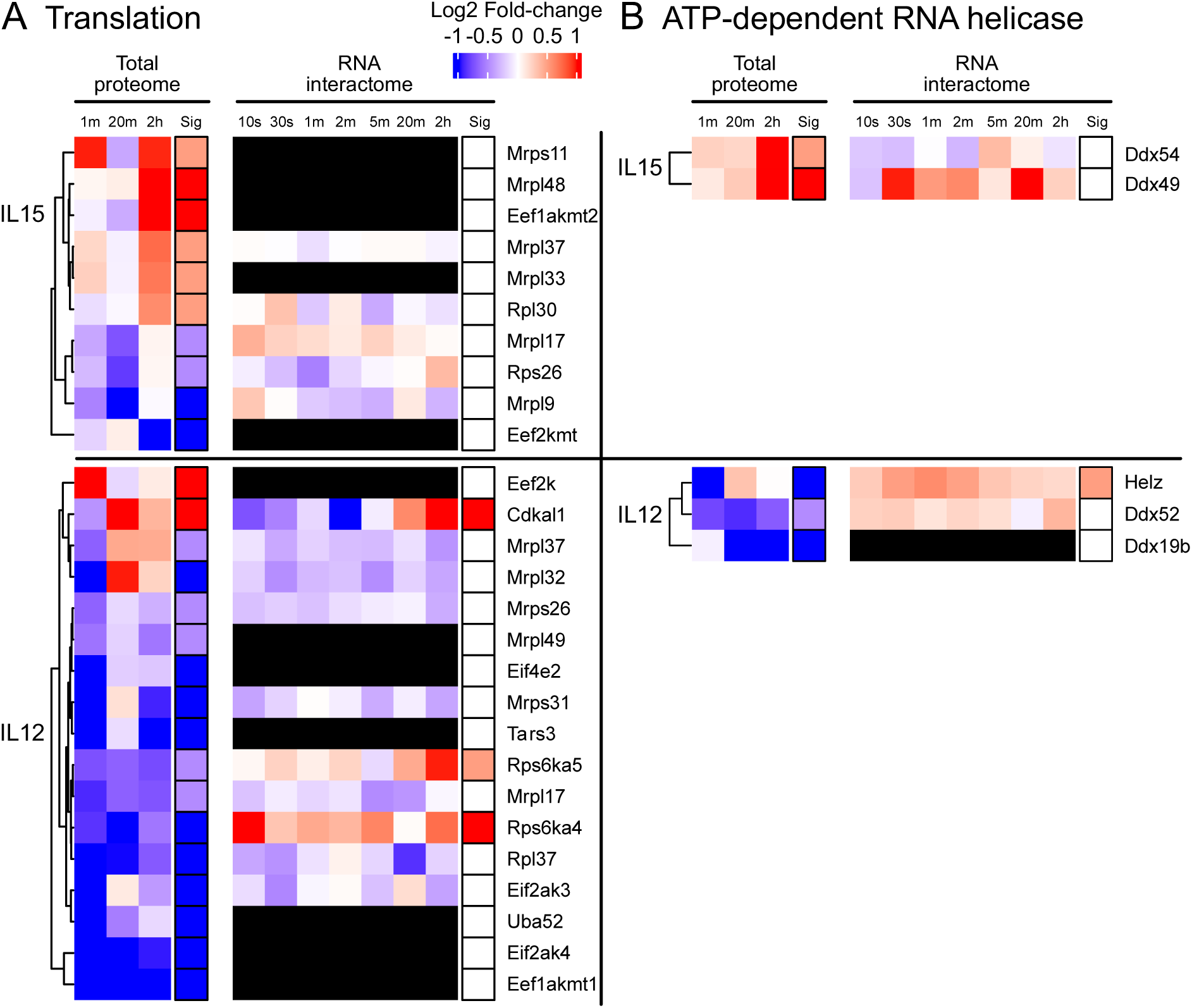
mRNA translation machinery and regulators significantly changed in cellular abundance following glucose withdrawal. Heatmap showing log2 fold change, from Glu+ to Glu-condition, in IL15 and IL12 cells, across experimental time course. Clustering was done based on the total proteome dataset. Black cells: protein not found in dataset. Statistical significance was determined by the lowest p value in the time course (two-tailed T Test, P<0.05), and the fold change at the corresponding time point (absFC>1.5: pale red/blue; absFC>2: full red/blue). (A) mRNA translation regulators were defined as experimentally validated components of GO:0006412 (Biological Process:Translation), plus all ribosomal proteins, translation initiation and elongation factors. (B) RNA helicases were defined as components of GO:0003724 (Molecular Function: ATP-dependent RNA helicase activity).

In IL15 cells eEF1AKMT2 was increased, and eEF2KMT decreased, at 2 hr. This suggests reduced global mRNA translation through impaired eEF2 methylation. In IL12 cells, eEF1AKMT1 was dramatically decreased across all time points, with a transient acute increase in eEF2K at 1 min. Decreased levels of eIF2α kinases, eIF2AK3 at 1 min, and eIF2AK4 at 20 min, were also observed. Together a mixture of predicted enhanced and repressed global translation was seen concurrently in IL12 cells.

Translation initiation, likely of specific mRNA subsets, may also be altered in IL12 cells. Decreased abundance of the initiation factor eIF4E2 at 1 min may indicate an acute and transient translation enhancement/inhibition of specific mRNAs, depending on its binding partners (GIGYF2 ^47^, 4E-T ^48^, miRNA ^49^, or TTP ^50^). IL12 cells also showed reduced abundance of the translation-promoting RNA-DNA helicase HELZ at 1 min, the stress-activated kinase RPS6KA4 (MSK2) at 20 min, the RNA-helicase DDX19B at 20 min, and the threonyl-tRNA synthetase TARS3 at 2 hr; the latter two being required for stress granule assembly.

Changes observed in IL12 cells indicated a transient initial halt in mRNA translation mediated by eEF2K and possibly eIF4E2, followed by a mixture of increased and decreased transcript-specific translation, supporting an adaptive response to the low glucose stress. Regulation at the level of transcript-specific translation has been previously reported during early T cell activation and effector stage ^51,52^. We speculate that related mechanisms are at play here, given the many shared characteristics between T cell activation and integrative stress response ^53^.

### Transcription factors can show opposing changes in cellular abundance versus RNA association

Transcription factors (TFs) are critical regulators of T cell activation, differentiation and metabolic adaptation, and were analyzed in detail. A list of TFs was obtained from a recent systems-level study in T cells ^54^, and analyzed in our datasets. The results are shown in Fig 4, with proteins changed in cellular abundance shown in Fig 4A, and those changed in RNA association shown in Fig 4B. Glycolytic IL12 cells had a greater number of TFs with significantly changed total abundance than IL15 cells following glucose depletion. Notably, comparing the cellular abundance and RNA-association datasets for TFs, we noted a dominant pattern, in which most altered TFs showed reduced total abundance, whereas increased RNA interactions by TFs predominated. We speculate that this de-coupled pattern reflects transient protection by RNA-TF association, potentially preventing these TFs from immediate degradation.

**Fig 4.**
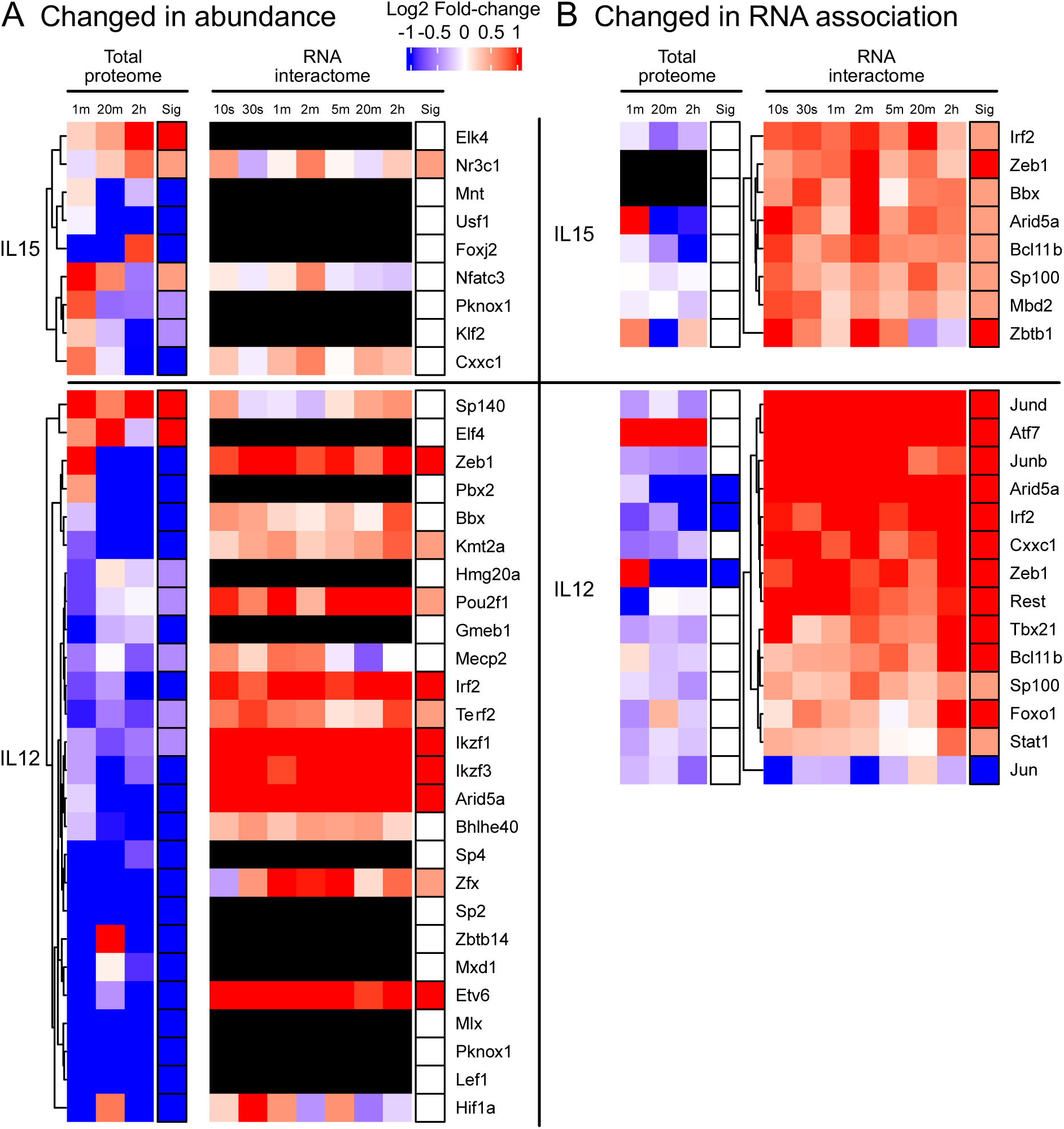
Key immunological transcription factors significantly changed in cellular abundance or RNA association following glucose withdrawal. Heatmap showing log2 fold change, from Glu+ to Glu-condition, in IL15 and IL12 cells, across experimental time course. Black cells: protein not found in dataset. Statistical significance was determined by the lowest p value in the time course (two-tailed T Test, P<0.05), and the fold change at the corresponding time point (absFC>1.5: pale red/blue; absFC>2: full red/blue). (A) Proteins changed in total cellular abundance, selected and clustered according to the total proteomic dataset. (B) Protein changed in RNA association, selected and clustered using the RNA-interactome dataset.

**Fig 5.**
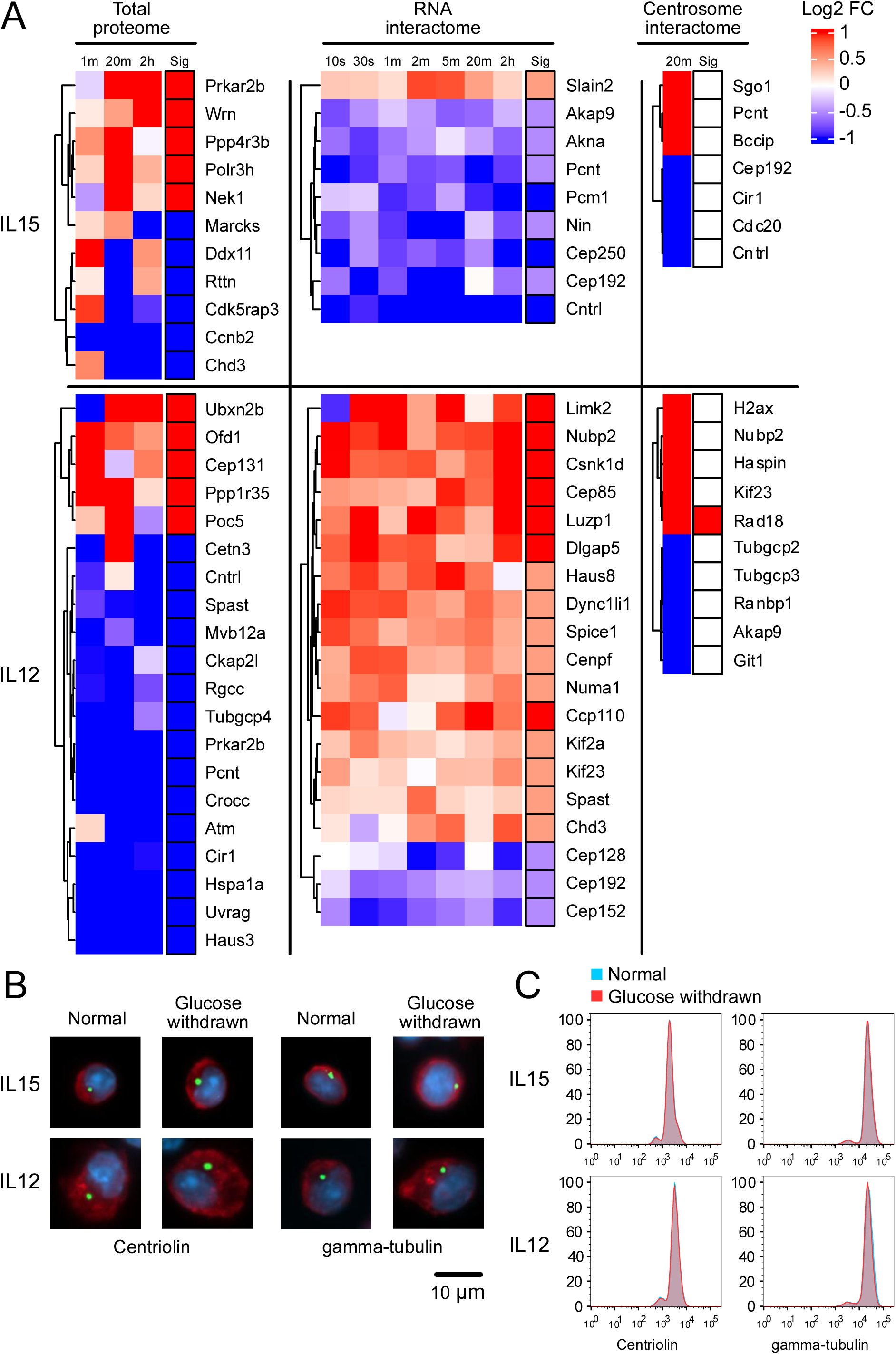
Altered RNA-centrosome association in response to glucose withdrawal in IL15 cells. (A) Heatmap showing log2 fold change, from Glu+ to Glu-condition, in IL15 and IL12 cells, across experimental time course. Significantly changed proteins in total proteome, RNA-interactome, and centrosome-associated proteome are shown. All indicated proteins are experimentally validated components of GO: 0005813 (Cellular Component: centrosome), and were selected according to their lowest p value (in total proteome and RNA-interactome) or fold-change (centrosome-interactome). (B) Immunofluorescence images of IL15 and IL12 cells in Glu+ or Glu-medium, using markers for structural components of centrosome (Centriolin and gamma-tubulin, green), actin cytoskeleton (Phalloidin, red), and nucleus (DAPI, blue). (C) Flow cytometry histogram showing intracellular Centriolin and gamma-tubulin intensities in Glu+ (blue) and Glu-(red) conditions.

In IL12 cells, several changes in total abundance were detected within one minute of glucose withdrawal, suggesting very active acute responses. Two factors were significantly increased: Type 1 interferon-dependent co-inhibitory receptor regulator SP140 ^55^ at 1 min and 2 hr, and T cell migration and differentiation factor ELF4 ^56^ at 20 min. In contrast, many more TFs were reduced, and only consistently decreased proteins are described below. The PBX-binding, T cell developmental factor PKNOX1 ^57^ and the PKNOX/PBX-binding proliferation regulator SP2 ^58^ were both decreased at all time points. PBX2, which may function as a complex with PKNOX1 and SP2, failed to be detected in Glu-condition at 20 min or 2 hr. The hypoxia-induced, glycolysis-promoting, T cell effector function regulator HIF1A ^59^ was also absent in all Glu-samples. Glucose-sensing glycolysis modulator MLX ^60^, T cell identity and exhaustion regulator LEF1 ^61^, and homeostatic proliferation factor ZFX ^62^ were also rapidly and persistently decreased. It is notable that HIF1A and MLX were specifically absent from IL15 cells, even on glucose, suggesting that these factors facilitate IL12 cell-specific responses.

In IL15 cells, TFs with increased abundance included the calcineurin-dependent T cell activation and proliferation factor NFATC3 ^63^ at 1 min, and ERK-dependent T cell differentiation factor ELK4 ^64^ at 20 min. Decreased TFs include T cell activation factor USF1 ^65^ at 20 min, MYC-antagonist anti-proliferation factor MNT ^66^ at 20 min, and activation and differentiation epigenetic regulator CXXC1 ^67^ at 2 hr.

Changes in TF-RNA association were relatively mild in IL15 cells, with most significant alterations seen in the first 2 min. These include interferon-induced T cell exhaustion factor IRF2 ^68^, pro-inflammatory cytokine-inducing factor ARID5A ^69^, memory T cell survival and homeostasis factor ZEB1 ^70^ and the T cell effector function and differentiation regulator BCL11B ^71^. These factors were also significantly changed in IL12 cells, suggesting a pan-T cell TF-RNA association in response to low glucose.

TFs in IL12 cells largely showed an immediate and persistent increase in RNA-association. These included the anti-apoptotic, proliferation and effector function regulator JUNB and JUND ^72–74^, longevity-associated epigenetic regulator ATF7 ^75^, and pro-inflammatory factor ARID5A, which showed consistently increased RNA interactions throughout the time course. Exhaustion factor IRF2, activation and differentiation epigenetic regulator CXXC1, survival and homeostasis factor ZEB1, and the epigenetic silencer and immune checkpoint regulator REST ^76^ also had increased associations, albeit less consistent. T cell migration, effector function and differentiation regulators TBET ^77^ and BCL11B ^71^, and naïve/memory T cell differentiation and survival factor FOXO1 ^78^ had increased associations mainly at 2 hr.

We note that Adenine-thymine (AT)-rich interactive domain containing protein 5a (ARID5A), was consistently increased in RNA-association under low glucose. ARID5A stabilizes *Tbx21* (encoding TBET) ^69^ and *Stat3* ^79^ mRNAs and may contribute to an inflamed/exhausted cellular state. For other consistently altered TFs, such as JUND and ATF7, there is limited prior evidence of RNA binding. It could be envisaged that they alter the stability of bound (m)RNAs, similar to ARID5A, or conversely RNA association might stabilize these proteins.

### Centrosome-RNA dissociation in IL15 cells

RNA-centrosome association has been postulated to facilitate centrosome assembly, microtubule nucleation and/or RNA localization ^80^. Individual mRNA transcripts identified as centrosome-associated by single molecule in-situ hybridization ^81^, mostly encode proteins essential for centrosome/cilial structure and functions. This suggests that RNA-centrosome association promotes site-specific mRNA translation ^82^. The centrosome plays an active role in T cells: in naïve or resting T cells the centrosome is located next to the nucleus; during T cell migration it is located at the uropod at the back of the moving cells; upon contact with target cells it is rapidly relocated to the site of contact, where it facilitates formation of the immunological synapse ^83^. The centrosome serves as an anchor for microtubule dynamics and intracellular active molecule delivery, critical for the movement and immunological function of T cells. A protocol for isolating centrosomal complexes (CAPTure) has been developed for human and mouse cells ^84^, and was used here to identify the centrosome-associated proteome. However, when this protocol was applied to identify RNA, the pulldown sequence dataset resembled the total cellular transcriptome, and was therefore omitted from further analyses. Since changes in RNA association with centrosomal proteins were most visible in the 2-20 min period (Table 1), we produced a centrosome-interactome dataset at the 20 min time point. Total proteome, RNA-interactome and centrosome-interactome datasets were compared, with a focus on experimentally validated centrosomal components (GO: 0005813 - Cellular Component: centrosome) (Fig 4A).

Some known centrosome interacting proteins changed in cellular abundance at 20 min in IL15 cells. Elevated components included centrosome integrity factor NEK1 ^85^, Cdk1/Cyclin B regulator PPP4R3B ^86^, and RNA polymerase III subunit POLR3H. Decreased proteins included centriole biogenesis and spindle pole integrity factor RTTN ^87^, and cell cycle and mTOR regulator CDK5RAP3 ^88^.

Although the RNA-protein interactome in IL15 cells showed few changes overall, these included key centrosomal structural proteins (CNTRL, PCNT, PCM1, and CEP250), for which reduced RNA-association was seen only in IL15 cells. CNTRL is essential for anchoring microtubules to centrioles, and required for cytokinesis and cell division ^89^. PCM1, together with pericentrin (PCNT), centrin, and ninein, form the pericentriolar satellites which are assembly sites for microtubules and transport hubs for centrosome-directed cargos ^90^. CEP250 (also known as C-Nap1) is a linker protein connecting the mother and daughter centrioles, preventing their premature separation ^91^. None of these proteins were altered in total cellular abundance.

Our CAPTure-MS centrosome pulldown dataset in LIL15 cells indicated that dissociation from RNA by these proteins was not accompanied by their exclusion from the centrosome. Furthermore, immunofluorescence visualization and flow cytometric quantitation of centrosomal markers (CNTRL and gamma-tubulin) confirmed that centrosomes remained intact after 20 min of glucose depletion (Fig 4B and C).

In IL12 cells changes in total abundance for centrosomal proteins were more prominent compared to the RNA-interactome and CAPture datasets. Increased abundance was seen for the mitotic progression and spindle position regulator UBXN2B ^92^ (only detected in Glu-condition) and the glycolysis-promoting mitotic spindle assembly factor SAC3D1 ^93^. Abundances were decreased for spindle microtubule-kinetochore attachment factor KNSTRN ^94^, mitotic spindle assembly checkpoint regulator ATM ^95^, pericentriolar material structural protein PCNT ^90^, cyclic AMP-dependent PKA regulator PRKAR2B ^96^, and the γ-tubulin ring complex component spindle pole formation factor TUBGCP4 ^97^.

In IL12 cells, changes in RNA interactions were mild and transient. Reduced RNA interaction was seen for scaffolding proteins CEP128, CEP152 and CEP192. Increased RNA association was seen for centrosome positioning protein CSNK1D ^98^, and other factors affecting centrosome integrity including CEP85, DLGAP5, LIMK2 and NUBP2. Proteins involved in ciliogenesis regulation, including LUZP1 ^99^ and CCP110 ^100^, also showed increased in RNA association. T cells do not form cilia, but immune synapses with target cells share many similar structures. LUZP1 and CCP110 potentially function in centrosome docking at the immune synapse during target cell encounter.

In the CAPture data for IL12 cells, centrosome association was unaltered for all of these proteins. Significantly changed centrosome binding was seen only for RAD18, an E3 ubiquitin-protein ligase involved in post-replicative DNA damage repair ^101^, suggesting the possibility of altered protein turnover.

## DISCUSSION

### Magnitude of RNA-Interactome reconfiguration depends on prior metabolic state

The ability of a cell to withstand nutrient fluctuation is intrinsically linked to its metabolic programming. Across evolution, many systems with high energy demands utilize glycolysis in preference to oxidative phosphorylation, including cytotoxic effector IL12 cells. This supports their very rapid proliferation and cytokine production following activation. However, while glycolysis is fast, it is also inefficient in ATP generation and requires a high flux of glucose. Effector T cells will quite frequently encounter low glucose environments, for example by actively migrating to sites of infection or tumor microenvironments. In contrast, memory IL15 cells predominately generate ATP through oxidative phosphorylation. They are long-lived and their primary function is to provide a persistent memory of the activation trigger, while being ready to transition back into effector cells upon antigen re-encounter. Presumably as a consequence, these two cell types show very different dynamics in RNA-protein interactions in response to low glucose challenge. IL12 cells showed substantial and rapid changes, likely actively adapting to the new environment; while in IL15 cells, the responses were relatively muted, with slower and fewer changes, perhaps prioritizing stability and long-term survival. Since effector IL12 CD8+ T cells more frequently encounter acute glucose deprivation, it might be expected that they would more actively respond to the challenge. Notably no proteins showing changes in RNA-association exhibited corresponding directional changes in total cellular abundance, indicating that these observations reflect genuine dynamics in RNA-interaction rather than fluctuations in protein synthesis or degradation.

Specifically, IL12 cells exhibited a biphasic response to acute glucose withdrawal: 67 proteins showed peak changes in RNA-protein association within the first minute, followed by a second major wave involving 109 proteins at two hours. Changes in IL15 cells were more modest, with changes peaking at intermediate time points (2-20 min), in only 42 proteins, and the response largely subsiding by two hours.

We propose that this distinction in low glucose response reflects the prior metabolic state of these cells. In principle, reliance on glycolysis requires a constant flow of glucose, and might make effector cells more susceptible to the effects of glucose withdrawal. However, we predict that the IL12 cell responses are protective and adaptive, making them better equipped at responding to glucose depletion. Environmental glucose level may serve as a “switch” for effector cells, notifying the cells that they have arrived at the site where their cytotoxic activity should commence. The oxidative memory cells do not have the role of cell killing at infection or tumour sites, and may therefore not show this specific response. This discrepancy between cell states was also reflected by changes in their total proteome, with 376/7131 proteins reduced in IL12 cells at immediate time point (1 min), compared to 13/6532 proteins in IL15 cells, suggesting an active macromolecular change specifically in IL12 cells.

Changes in RNA-binding by mRNA translation regulators were exclusively seen in IL12 T cells; notably reduced RPS5 association in the immediate response, followed by reduced interactions for RPS3 and eIF4B at later stages. A larger number of translation regulators changed at the level of total cellular abundance, likely caused by active protein degradation. Notably, factors altered in RNA association did not overlap with factors changed in cellular abundance, suggesting independent protein regulation occurring at multiple levels during the low-glucose response.

In budding yeast, translation inhibition following acute glucose deprivation has been linked to extremely rapid ATP depletion, from ∼2 mM to ∼150 µM over ∼30 sec. This prompts a global arrest of translation through displacement of DEAD-box helicase initiation factors, eIF4A and Ded1, from translation initiation regions of mRNAs. These have Km values for ATP binding of around 300-500 µM ^102,103^ and are required for eIF4B binding at the translation start site. Quantitation of intracellular NTP levels in IL12 T cells revealed that ATP levels are also rapidly depleted following glucose withdrawal, from ∼2.4 mM to ∼1.5 mM in the first minute. However, the degree of ATP depletion is less profound and would not be expected to cause the dissociation of translation regulators. Consistent with this, eIF4A and the RNA helicase DDX3X (the mammalian homolog of yeast Ded1) remained stably associated with RNA in T cells. Moreover, the reduction seen for eIF4B-RNA interactions in IL12 cells was considerably slower than in yeast, suggesting a different mechanism. We conclude that the RNA-interactome reconfiguration in glycolytic IL12 T cells is not a consequence of a global bioenergetic switch, but rather an active signaling adaptation to metabolic stress.

### Transcription Factors as RNA-Binding Proteins

An unexpected finding was the rapid and significant alteration in RNA-interactions observed for multiple transcription factors following glucose withdrawal. In the immediate response (1 min), both cell states showed increased RNA-association for transcription factors critical for survival, including the AP-1 complex components JUNB and JUND. Both JUNB and JUND are anti-apoptotic, with JUNB notably promoting glycolysis and controlling the expression of cytotoxic molecules including IFNγ. We speculate that rapid binding to RNA could reduce the availability of these transcription factors for active transcription of energy-demanding genes at the chromatin level. Consistent with this hypothesis, most transcription factors that showed altered total cellular abundance were depleted. At least in 8 out of 24 instances, the reduction occurred at 1 min and persisted throughout the time course, again suggesting rapid, active targeted protein degradation. We postulate that specific degradation helps commit the cells to a new functional state, while factors sequestered by RNA remain available for use once the stress response is completed. This may represent a non-canonical layer of stress regulation.

We note that several RNA-interacting transcription factors are established regulators of cellular stress responses. For instance, SIRT1, which showed increased RNA binding in IL12 cells at 2 hours, is a key metabolic regulator that restrains glycolysis and interacts with c-JUN and TBET to orchestrate T cell differentiation. ARID5A, which demonstrated increased RNA association in both cell types, is known to stabilize *Tbx21* (TBET) and *Stat3* mRNA, preventing their degradation and contributing to an inflamed or exhausted cellular state. The engagement of these factors with qRNA suggests a post-transcriptional regulatory network. This may affect the stability of both RNAs and proteins, during the functional switch in IL12 T cells. The molecular consequences of these altered RNA-associations will form the basis for future, more focused studies.

### Divergent Centrosomal RNA Dynamics

The centrosome has specialized roles in T cells, functioning as a critical anchor for microtubule dynamics, cellular migration, and the formation of the immunological synapse upon target encounter. Recent evidence indicates that mRNAs physically localize to the centrosome to facilitate the site-specific translation of proteins vital for maintaining these functions. Several centrosomal proteins were identified in our datasets as RNA-binding: 41 in IL15 cells and 68 in IL12 cells. A striking divergence between the cell states was the specific reduction in RNA-centrosome interactions in IL15 cells. At 20 min post-glucose withdrawal, IL15 cells exhibited reduced RNA association with several core, structural components of the centrosome, including CNTRL, PCNT, PCM1, and CEP250.

The reduced recovery of these proteins with RNA in IL15 cells was not accompanied by alterations in their cellular abundance or centrosomal association. Nor did the centrosome physically disassemble, as immunofluorescence confirmed that centrosomes remained structurally intact. This suggests a “regulatory pause” within memory cells, possibly reflecting selective inhibition of localized translation at the centrosome. This might help memory cells conserve metabolic resources, to better survive under a low resource environment. In contrast, IL12 cells displayed no strong reduction in interactions between RNA and core centrosomal proteins. Indeed, many RNA interactions were increased. IL12 cells also showed increased centrosome binding by RAD18, an E3 ubiquitin ligase which may participate in localized protein turnover. This again indicates that sudden glucose withdrawal in active effector cells triggers active change, rather than a conservative pause state.

## Conclusion

Reconfiguration of the RNA-interactome under glucose depletion stress does not elicit a uniform cellular response, but specific programs defined by immunological differentiation. Glycolytic effector cells mount a strong, biphasic response involving the active engagement of stress-responsive transcription factors and RNA helicases, likely adapted to environments where they need to function. In contrast, OxPhos-reliant memory cells exhibit a muted response, but this included uncoupling of RNA from centromeric proteins. These findings emphasize that metabolic modality does not merely fuel the cell but defines how the cell perceives and survives environmental stress.

## RESOURCE AVAILABILITY

Requests for further information and resources should be directed to and will be fulfilled by the lead contact, Thomas Tan.

The mass spectrometry proteomics data have been deposited to the ProteomeXchange Consortium via the PRIDE (https://www.ebi.ac.uk/pride/) partner repository with the dataset identifier PXD079329. Analyzed Excel files and codes used in R are available at Open Science Framework, https://osf.io/ (DOI: 10.17605/OSF.IO/N3RM7).

## ACKNOWLEDGMENTS

The authors thank Prof. Amy Buck for her provision of primary cell culture facilities and access to flow cytometer; Prof. Rose Zamoyska and Prof. Val Brunton for providing experimental mice; Prof. Andrei Chabes for NTP quantitation. Heatmaps in all figures were made using ComplexHeatmap (ver. 2.24.1) ^104^. Funding for this study was provided by Wellcome Principal Research Fellowship [222516 to D.T.].

## AUTHOR CONTRIBUTIONS

Conceptualization, T.T and D.T.; Data curation, T.T. and C.S.; Formal analysis, T.T.; Funding acquisition, D.T.; Investigation, T.T.; Methodology, T.T. and C.S.; Project administration, D.T.; Resources, T.T.; Software, T.T.; Supervision, D.T.; Validation, T.T.; Visualization, T.T.; Writing – original draft, T.T.; Writing – review & editing, D.T.

## STAR Methods

### Experimental model

Mice used in experiments were of C57BL/6 background strain carrying homozygous OT-1 transgenic T cell receptor on a Rag-1KO background. All mice were bred at the animal facility in the University of Edinburgh in accordance with the UK Home Office and local ethically approved guidelines. Mice of both sexes were used.

### Cell culture and glucose withdrawal

Primary T cells were isolated ex vivo from lymph nodes of 6-8 weeks old mice. Cells were activated with 100 nM cognate SIINFEKL peptide (custom synthesized by Cambridge Peptides) for two days, in fresh IMDM medium (Sigma-Aldrich, I3390) supplemented with 10% FCS (Sigma-Aldrich, F9665), 2 mM L-glutamine (Gibco, 25030081), 100 U/ml penicillin+streptomycin (Capricorn Scientific, CSR403) and 50 μM β-mercaptoethanol (Thermo Scientific, 125472500). Activated cells were washed free of the peptide, and transferred to a fresh culture medium containing 20 ng/mL human IL-2 cytokine (PeproTech, 200-02-1MG), in which the cells proliferate for two days. To induce differentiation, the cells were then either transferred to a fresh IL-2-containing SILAC heavy (containing 50 μg/L each of ^13^C_6_-L-lysine and ^13^C_6_-L-arginine, CK Isotope, CLM-2247-H and CLM-2265-H) or light (containing 50 μg/L each of L-lysine and L-arginine, Sigma-Aldrich, L9037 and A6969) medium (RPMI 1640, Thermo Scientific, A33823) and further cultured for two days, before 5 ng/mL mouse IL-12 (PeproTech, 210-12-10UG) was added and cultured for one day; or cultured in a fresh SILAC heavy/light medium containing 20 ng/mL human IL-15 (PeproTech, 200-15-1MG) for three days. On the day of harvest, Glu+ (glucose-containing, Thermo Scientific, A33823) or Glu-(glucose-deprived, Thermo Scientific, A2494201) SILAC heavy/light media containing the same cytokines were prepared, and pre-warmed at 37°C in a water bath to minimize temperature-related acute stress response. Cells were counted and 40 million cells were spun down and medium replaced with Glu+ or Glu-(conditional media) for set durations (10s, 30s, 1m, 2m, 5m, 20m, 2h). For 20m and 2h, cells were incubated at 37°C in 5mL conditional medium until 7 min before the set duration, then spun for 5 min, and transferred in 1 mL conditional medium to a 10 cm dish pre-incubated with the culture medium. Cells were crosslinked with UVC, twice with 300 mJ/cm2 each, with cells mixed by gentle shaking in between. Crosslinked cells were collected in 2 mL microcentrifuge tubes, spun down and pellet frozen on dry ice. For 10s, 30s, 1m, 2m, and 5m time points, cells were resuspended directly into 1 mL condition media and incubated directly on dishes, and were crosslinked at precise times. Samples were stored at −80°C until RNP extraction. For total proteomes and CAPture centrosome-pulldown proteomes, cell culture and glucose withdrawal were performed using IMDM medium. For CAPture experiment, 0.1 mg/mL cycloheximide was added to Glu+ or Glu-medium for the last 10 min of culture, with no UV-crosslinking before harvesting.

### TRAPP RNA-interactome

Total cellular ribonucleoprotein was extracted from 30 million cells (IL12) or 60 million cells (IL15), with 1.5mL Trizol. Lysate was passed 6 times through an insulin needle, heated to 80°C for 5 min, and spun at 16,000 rcf for 10 min at 4°C. Supernatant was collected and 50 µL was extracted for RNA, and quantitated using a Quant-iT Broad Range RNA Assay Kit (Thermo Scientific, Q10213). For each IL15 replicate, 50 µg RNA equivalent of light lysate was pooled with 50 µg of heavy lysate; while for each IL12 replicate, 90 µg of light/heavy lysates were pooled.

The pooled lysate was loaded onto two EconoSpin columns for RNA (Epoch Life Science, 1940–250) and each column washed once with 600 μL WB1A (3 M guanidine thiocynate, 1 M sodium acetate pH 4.0, 30% ethanol). Collection tubes were replaced, and the columns were washed once with 600 μL WB1B (2 M guanidine hydrochloride, 20 mM Tris–HCl pH 6.4, 50% acetone, 20% ethanol), twice with 600 μL WB2B (20 mM Tris–HCl pH 6.8, 0.5 mM EDTA, 80% ethanol) and then once with 500 μL WB3B-80 (80% acetonitrile, 50 mM Tris–HCl pH 7.8, 2 mM CaCl 2). After the last wash columns were spun for 2 min at 16,000 rcf. Collection tubes were replaced with 1.5 mL low protein binding tubes (Thermo Scientific, 90410). To each column was added 0.25 μg Trypsin + Lys-C (Promega, V5073) in 75 μL WB3B-60 (60% acetonitrile, 50 mM Tris–HCl pH 7.8, 2 mM CaCl 2) with incubation for 16 h at 37 °C. The two columns were eluted by spinning at 16,000 rcf for 30 sec, and each column was washed with 75 μL WB3B-60. Eluates from the two columns were pooled and spun at 16,000 rcf for 5 min. Supernatant was transferred to a new tube. 1M DTT (Sigma-Aldrich, 43816) was added to the eluate to a final concentration of 5 mM and the mixture was incubated at 37 °C for 20 min. Peptide alkylation was done with 10 mM final Iodoacetamide (IAA, Merck, A3221) at RT in the dark for 20 min, followed by inactivation with 10 mM final DTT, incubated at RT for 5 min. 0.15 μg Trypsin + Lys-C in 7.5 μL WB3B-60 was added and incubated at 37 °C for 1 h for complete digestion of the peptides. The volume of the peptides was evaporated to ∼10 μL and resuspended in 100 µL 0.1% Trifluoroacetic acid (TFA, Thermo Scientific, 85183), acidified with 12 µL 10% TFA, and desalted with C18 stage tips. Mass spectrometry data was acquired as single-shot DIA SILAC and analyzed in DIA-NN (ver. 1.8.1) using a gas-phase fractionated DIA spectral library constructed using pooled Glu+ IL15 and IL12 samples.

RNA-interactome data was analyzed using the workflow described in Fig. S1.

### Total proteome

Total proteins were extracted from 20 million cells, with 350µL RIPA Buffer (Enzo Life Sciences, SV-39244-02). Lysate was sonicated (Diagenode Bioruptor Pico, 3 cycles, 30sec on, 30sec off) and spun at 15,000 rcf for 10 minutes at 4°C, supernatant was collected. Lysate was quantitated using Quant-iT Qubit Protein Assay Kit (Thermo Scientific, Q33212). 100 ug of total protein was taken from each sample, topped up to 100 µL with 100 mM Triethylammonium bicarbonate buffer (TEAB, Sigma-Aldrich, T7408) if necessary. Samples were reduced with 10 mM DTT and then alkalised with 18.75 mM IAA, then precipitated with 6X volume of acetone overnight. Protein was resuspended in 100 µL 50 mM TEAB, then digested with 2.5 µg Trypsin+LysC at 37°C overnight. On the next day an extra 0.5 µg Trypsin+LysC was added to the samples and incubated for another 1hr. Samples were then dried down and resuspended in 100 µL 0.1% TFA, acidified with 12 µL 10% TFA, and desalted with C18 stage tips. Mass spectrometry data was acquired as single-shot label-free DIA.

Total proteome data was analyzed using the workflow described in Fig. S2.

### CAPture centrosome-pulldown proteome

Total protein was extracted from 60 million cells, in 300 µL CAPture buffer (50 mM Tris-HCl pH 8.0, 300 mM NaCl, 0.2 %(v/v) Igepal CA-630, 10 %(v/v) glycerol, 0.1 mg/mL cycloheximide (APExBIO, A8244), protease inhibitor cocktail (Complete EDTA-free, Roche, 05892791001) and phosphatase inhibitor cocktail (PhosStop; Roche, 4906845001)). Lysate was sonicated briefly (Diagenode Bioruptor Pico, 30sec on/30sec off, 3 cycles at 4°C) and centrifuged at 1,800 rcf for 10 minutes at 4 °C. Immunoprecipitation was performed with the lysate for 2h at 4°C, with end-to-end rotation at 12 rpm, using Biotin-CAPture peptide (Biotin-SPSPTGGRALRFDPTAFVKAKERKQREIQMKQQ, 2BScientific, 6 µL of 5 mg/mL used) coupled with 60 µL streptavidin M280 dynabeads (Thermo Scientific, 11205D), or with dynabeads without the CAPture pepitde. Following 4 washes with 300 µL CAPture buffer, and 2 washes with 500 µL detergent-free CAPture buffer. Beads were resuspended in 300 µL WB3B-60. Samples were reduced with 10 mM DTT and then alkalized with 18.75 mM IAA, and digested using 0.3µg trypsin+Lys-C overnight. Digested peptides were dried to <10µL and resuspended in 150µL 0.1% TFA, then acidified with 10% TFA and desalted with C18 stage tips. Mass spectrometry data was acquired as single-shot label-free DIA.

CAPture proteome data was analyzed using the workflow described in Fig. S3.

### Immunofluorescence

Cells were fixed in Intracellular Staining Fixation Buffer (BioLegend, 420801) and permeabilized in Intracellular Staining Permeabilization Wash Buffer (BioLegend, 421002). Antibodies against centriolin (Santa Cruz, sc-365521) and gamma tubulin (Invitrogen, MA1-850) were used. Cells were incubated with primary antibodies for 16 hours at 4°C, washed, followed by staining with PE rat anti-mouse IgG1 (BioLegend, 406608) or IgG2b (BioLegend, 406708) secondary antibody for 1 hour at RT. After washes, actin filaments were then stained with Alexa Fluor® 647 Phalloidin (Cell Signaling Technology, 8940). Cells were washed and mounted on slide with ProLong™ Gold Antifade Mountant with DAPI (Thermo Fisher Scientific, P36931). Images were acquired using an Axio Observer light microscope (Zeiss).

### Flow cytometry

Cells stained for immunofluorescence, before staining with Phalloidin and DAPI, were transferred to FACS Buffer (0.5% BSA, 0.01% Sodium Azide, 2mM EDTA in PBS) and acquired on a MACSQuant Analyzer 10 flow cytometer (Miltenyi Biotec). All analyses were done in Flowjo 10, on intact singlet events.

### Cellular NTP quantitation

10 x 10^6^ cells were transferred into a 2 mL microcentrifuge tube and spun at 300 x g for 5 min. Supernatant were removed and cells were resuspended in 1 mL of pre-warmed Glu+ or Glu-conditional medium, and incubated at 37°C. Cells were spun down and lysed at precise time points (1m, 10m, and 2h). Cells were lysed by resuspending pellet in 1 mL TCA Buffer (12% Trichloroacetic acid (w/v), 15 mM MgCl_2_) on ice. The lysates were snap frozen in dry ice, and stored at −80°C. All time points and conditions were tested in biological triplicate.

For NTP quantification, thawed samples were vortexed for 30 sec then centrifuged at 4,000 rpm for 5 min at 4°C. The supernatant was transferred to a 1.5 ml Eppendorf tube and neutralized twice in a 1:1 ratio with a dichloromethane (DCM)–trioctylamine (TOC) mixture (1 ml DCM and 0.28 ml TOC). Following centrifugation at 14,000 rpm for 2 min at 4°C, 900 μL of the supernatant was transferred to new Eppendorf tubes. A 100 μL aliquot of the extracted nucleoside solution was then mixed with 1.15 mL Milli-Q water, and the pH was adjusted to between 3 and 4 using 0.5 μL 6 M HCl. Subsequently, analysis was performed by HPLC UV as previously described ^105^. NTP quantitation was normalized by total protein content in the same sample, and converted to mM per cell, assuming 686 fL of cellular water content (80% of total cellular volume) in effector CD8 T cells ^106,107^.

## RESOURCE IDENTIFICATION INITIATIVE

Antibodies: for immunofluorescence and flow cytometry, cells were stained with mouse monoclonal IgG1κ Centriolin antibody (Santa Cruz sc-365521, RRID: AB_10851483) and mouse monoclonal IgG2b gamma tubulin antibody (Invitrogen MA1-850, RRID: AB_2211249). Secondary antibodies used were PE rat anti-mouse IgG1 (BioLegend 406608, RRID: AB_10551618) or IgG2b (BioLegend 406708, RRID: AB_2563381).

---

**Fig S1.**
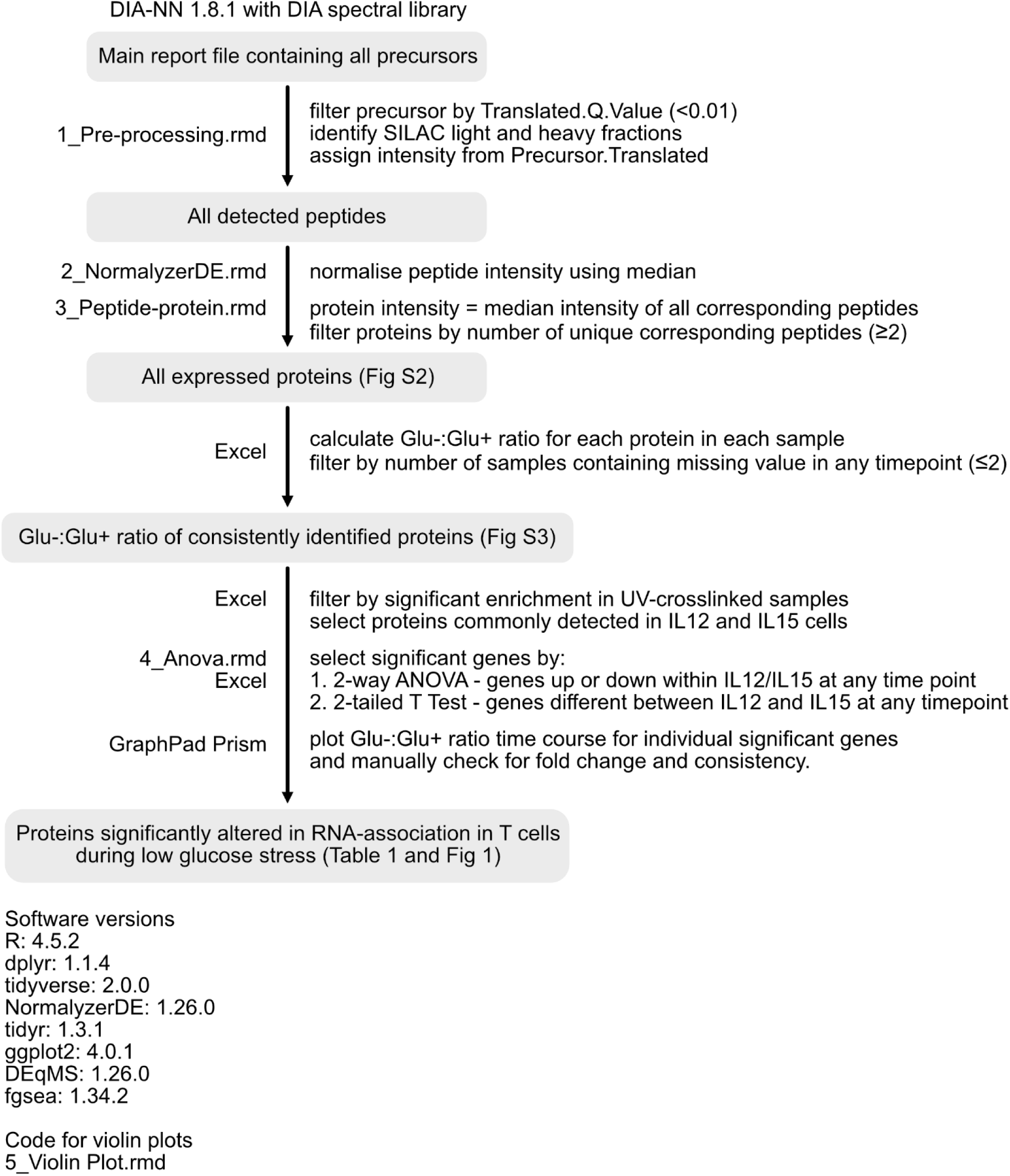
Analysis workflow for RNA-interactome.

**Fig S2.**
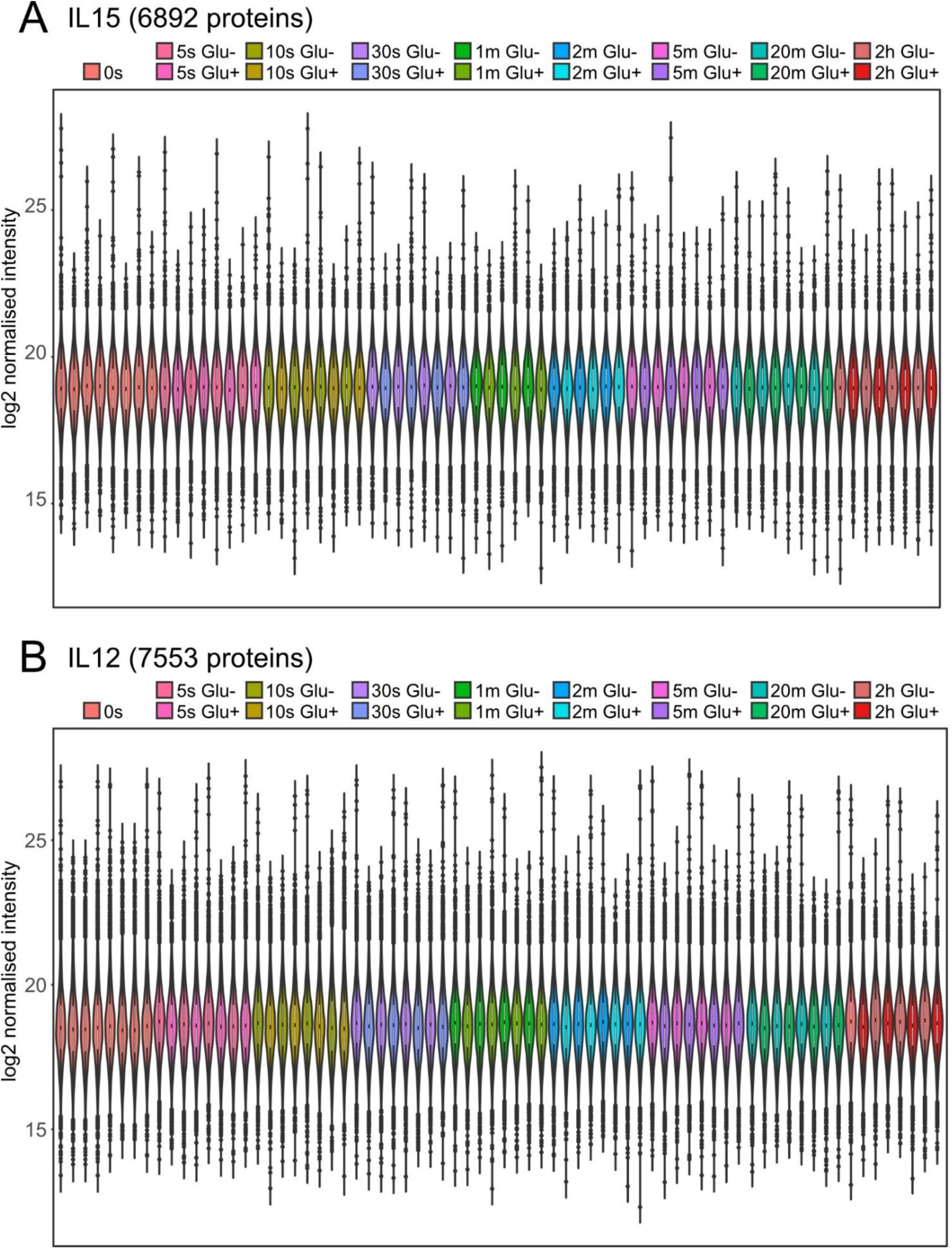
Median-normalised protein intensities in Glu+ and Glu-samples of IL15 and IL12 cells.

**Fig S3.**
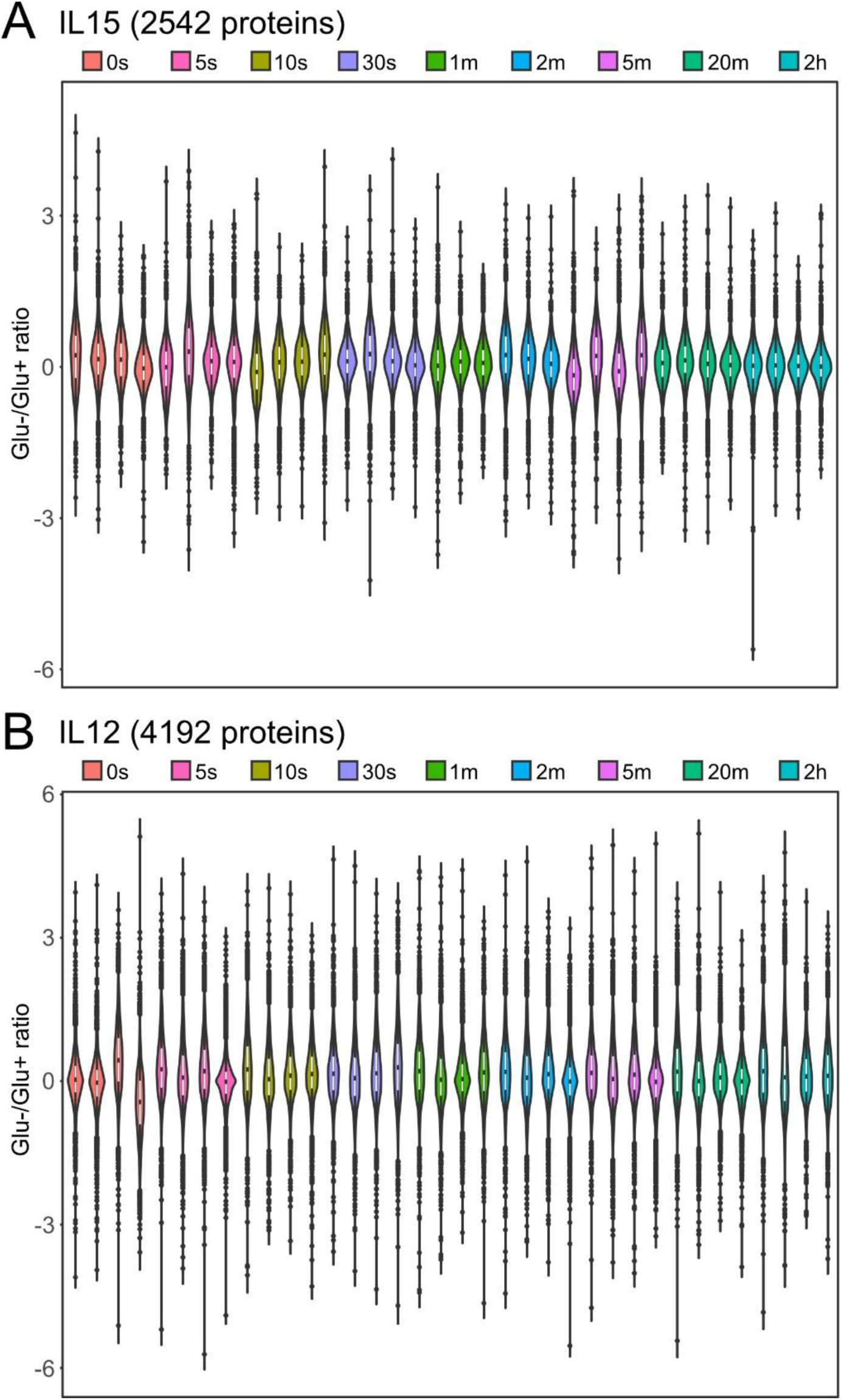
Glu-:Glu+ ratios of consistently identified proteins in IL15 and IL12 cells.

**Fig S4.**
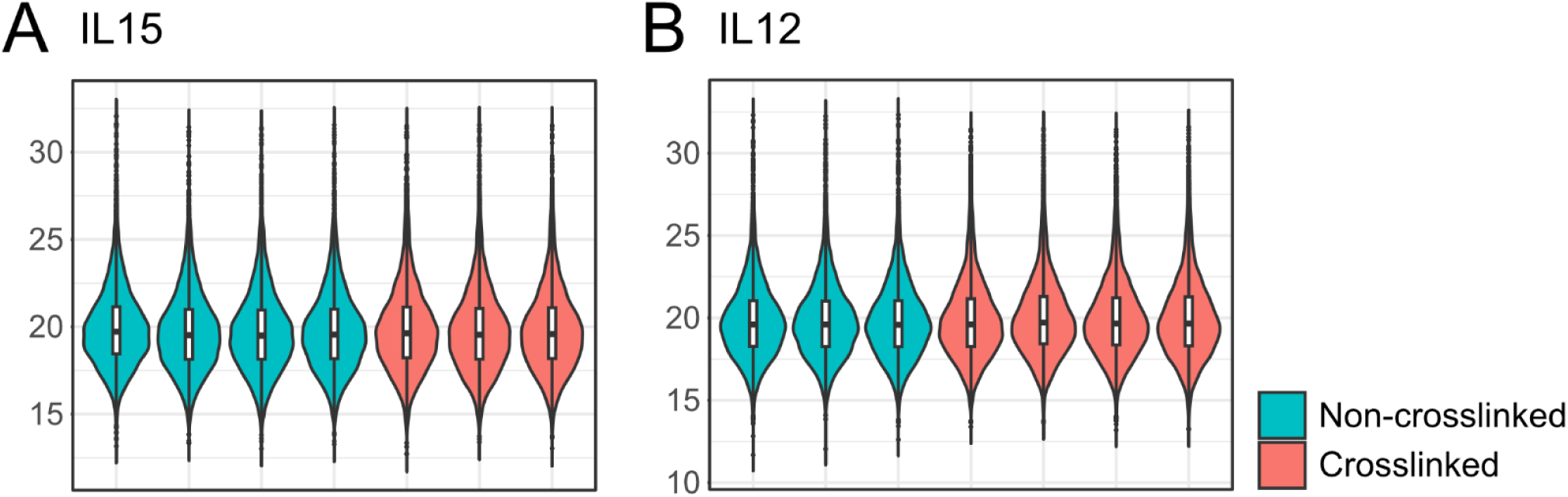
Vsn-normalised RNA-interactome protein intensities in IL15 and IL12 cells, with/without UV-crosslink.

**Fig S5.**
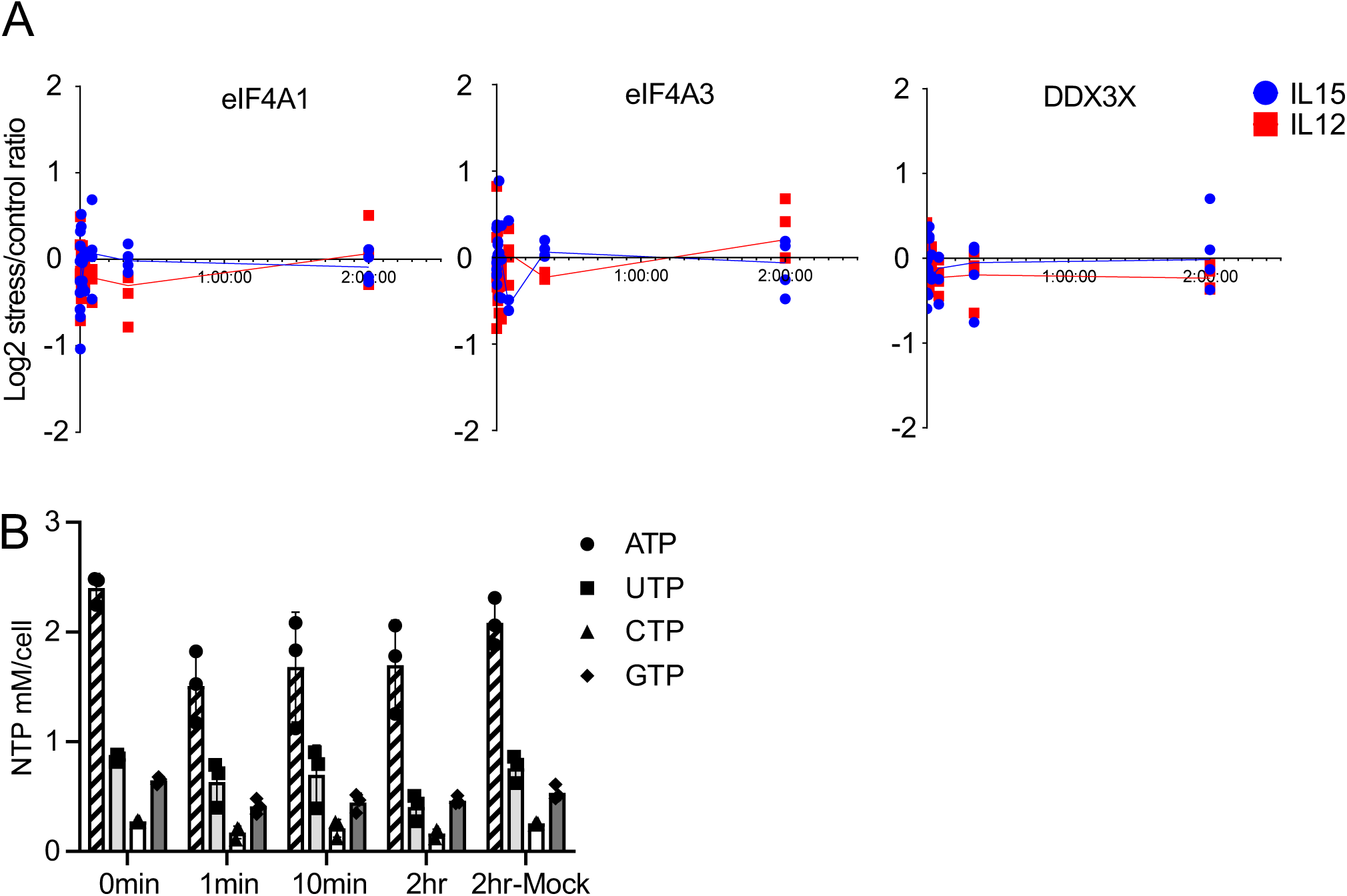
Quantitation of intracellular NTP levels at immediate (1 min), intermediate (10 min) and later (2 hr) time points of glucose withdrawal in IL12 cells. NTP quantitation was normalised in each sample by total protein quantitation, to control for loading. Quantity was measured as pmole per 10 million cells, and was converted to mM per cell, assuming 686 fL of cellular water content. Circular dots in textured bar: ATP; square dots in pale grey bar: UTP; triangular dots in white bar: CTP; diamond dots in dark grey bar: GTP. Error bar: S.D. of 3 biological samples.

**Fig S6.**
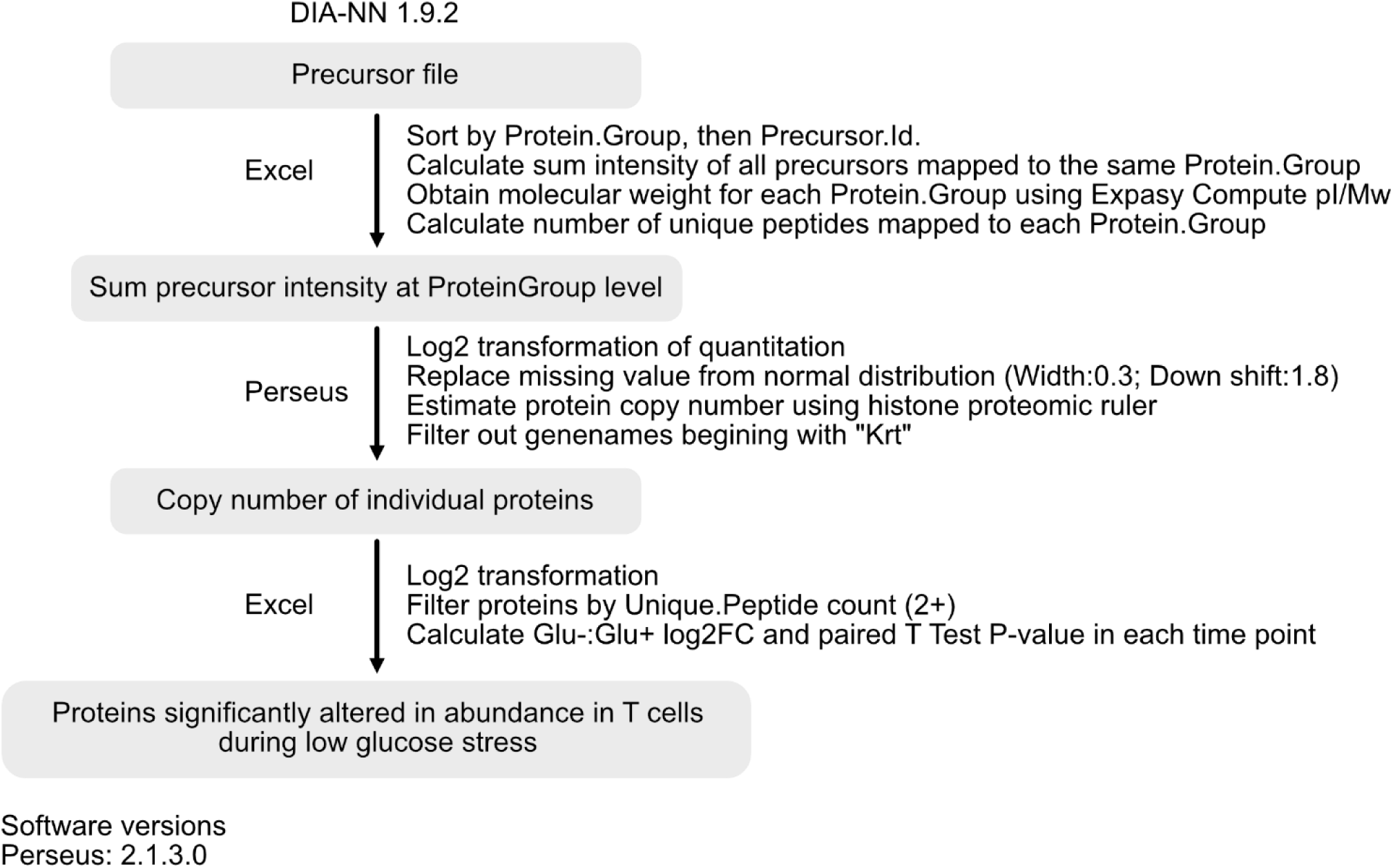
Analysis workflow for total proteome.

**Fig S7.**
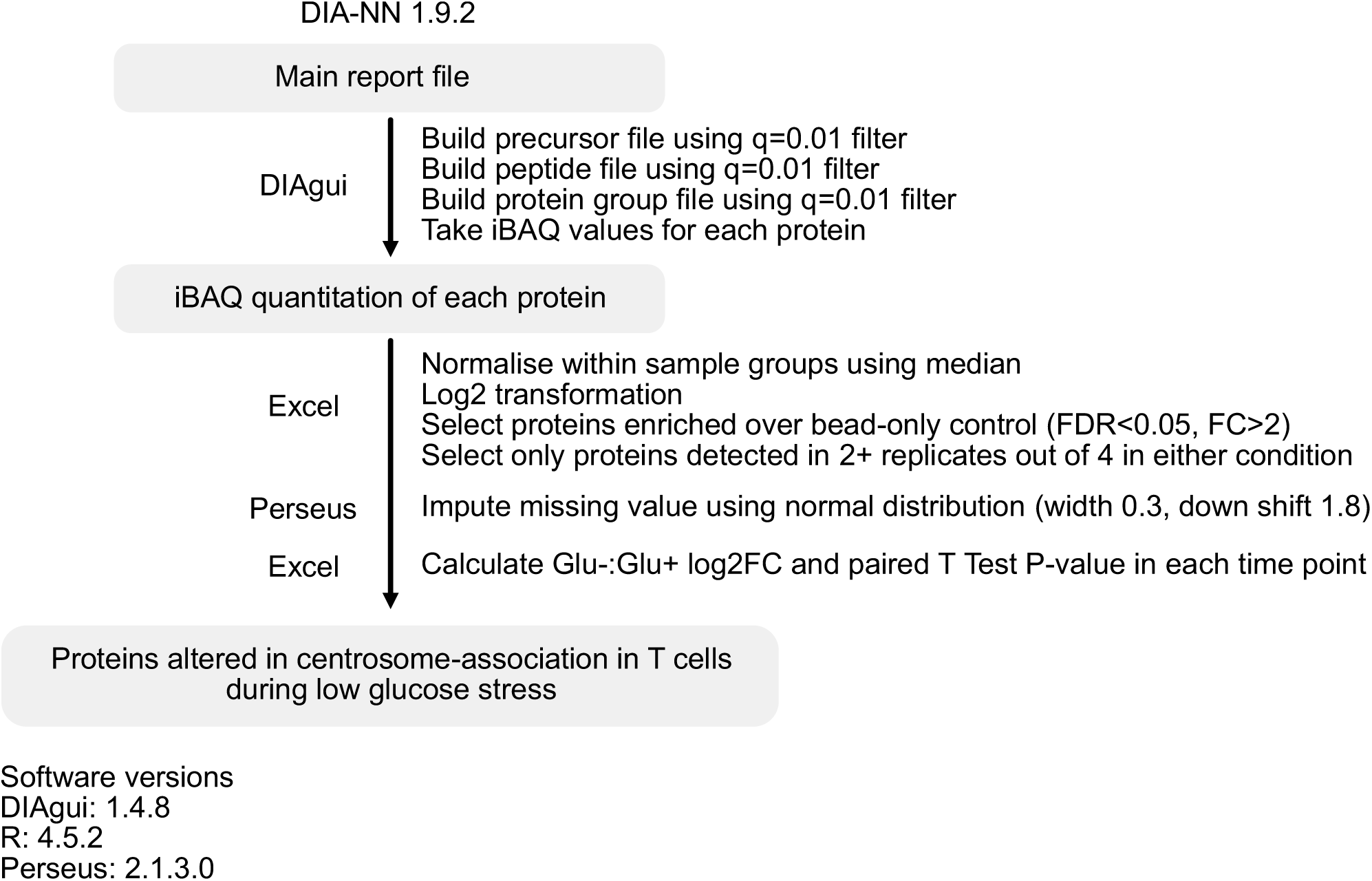
Analysis workflow for CAPture centrosome-associated proteome.

**Fig S8.**
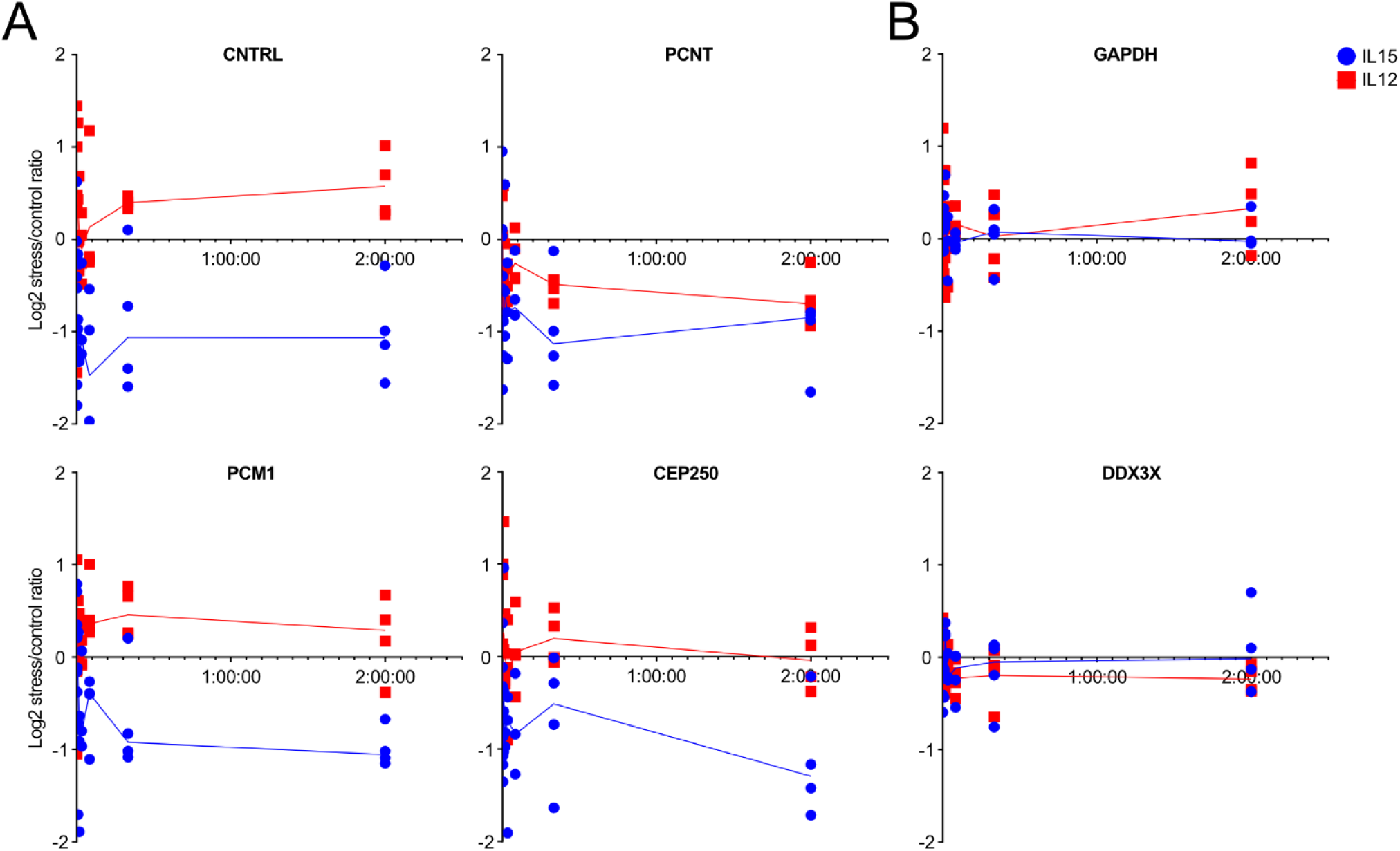
Changes in RNA-association of key centrosomal proteins across the glucose withdrawal time course. Stress/control (Glu-:Glu+) protein intensity ratio of (A) centrosomal proteins CNTRL, PCNT, PCM1 and CEP250; (B) examples of unchanged proteins GAPDH and DDX3X were plotted showing individual replicates in the RNA-interactome dataset. Blue dots: IL15; red dots: IL12 cells; Lines represent median of all replicates.

